# DYNLRB1 is Essential for Dynein Mediated Transport and Neuronal Survival

**DOI:** 10.1101/727016

**Authors:** Marco Terenzio, Agostina Di Pizio, Ida Rishal, Letizia Marvaldi, Pierluigi Di Matteo, Riki Kawaguchi, Giovanni Coppola, Giampietro Schiavo, Elizabeth M.C. Fisher, Mike Fainzilber

## Abstract

The cytoplasmic dynein motor complex transports essential signals and organelles from the cell periphery to perinuclear region, hence is critical for the survival and function of highly polarized cells such as neurons. Dynein Light Chain Roadblock-Type 1 (DYNLRB1) is thought to be an accessory subunit required for specific cargos, but here we show that it is essential for general dynein-mediated transport and sensory neuron survival. Homozygous *Dynlrb1* null mice are not viable and die during early embryonic development. Furthermore, heterozygous or adult knockdown animals display reduced neuronal growth, and selective depletion of *Dynlrb1* in proprioceptive neurons compromises their survival. Conditional depletion of *Dynlrb1* in sensory neurons causes deficits in several signaling pathways, including β-catenin subcellular localization, and a severe impairment in the axonal transport of both lysosomes and retrograde signaling endosomes. Hence, DYNLRB1 is an essential component of the dynein complex.

## INTRODUCTION

Molecular motors perform a critical function in eukaryotic cells, ensuring the anterograde (kinesins) and retrograde (dyneins) transport of cargoes, organelles and cellular components. Dynein motors are divided into two major classes: axonemal dyneins, which are involved in ciliary and flagellar movement, and cytoplasmic dyneins, minus-end microtubule molecular motors involved in vesicular transport, cell migration, cell division and maintenance of Golgi integrity [1–3]. Cytoplasmic dynein is the main retrograde motor in all eukaryotic cells and is particularly important in highly polarized cells such as neurons, where it carries essential signals and organelles from the periphery to the cell body [4]. Cytoplasmic dynein consists of two heavy chains (HC) associated with intermediate chains (IC), which provide direct binding to dynactin [5–7], four light intermediate chains (LIC) and several light chains (LC) [3]. There are three families of dynein light chains, LC7/road-block, LC8 and Tctx1/rp3, the latter playing a role in the direct binding of cargoes to the dynein motor complex [3,8,9].

Dynein Light Chain Roadblock-Type 1 (DYNLRB1), also known as Km23, and DNLC2A [10, 11], was first identified in Drosophila during a genetic screen for axonal transport mutants [12]. *Roadblock* mutants exhibited defects in intracellular transport, including intra-axonal accumulation of synaptic cargoes, severe axonal degeneration and aberrant chromosome segregation [12]. DYNLRB protein sequence is highly conserved among different species; with two isoforms, DYNLRB1 and DYNLRB2, sharing 98% sequence similarity in mammals [13]. Mammalian DYNLRB1 has been studied mainly as an adaptor linking specific modules to the dynein complex, including SMAD2 complexes activated by TGFβ receptors [11,14–18] and Rab6 or N-acetyl-D-glucosamine kinase (NAGK) for interactions with the Golgi compartment [19–21].

In previous work, we identified a role for the dynein complex in regulating axon growth rates [22, 23]. A screen of all dynein accessory subunits now shows that knockdown of *Dynlrb1* reduces neurite outgrowth in cultured sensory neurons. Surprisingly, a complete knockout of *Dynlrb1* was found to be embryonic lethal, suggesting the existence of non-redundant functions between DYNLRB1 and DYNLRB2. Conditional knockout or acute knockdown of *Dynlrb1* compromised survival of proprioceptive neurons, together with changes in subcellular localization of transported cargos, and major impairments in the axonal transport of lysosomes and retrograde signaling endosomes. Thus, DYNLRB1 is an essential component of the dynein complex, and the conditional allele described here enables selective targeting of dynein functions.

## RESULTS

### Depletion of *Dynlrb1* impairs neurite outgrowth in cultured DRG neurons

We performed an siRNA screen in cultured dorsal root ganglion (DRG) neurons to investigate the effects of downregulating individual dynein subunits on axonal growth. Knockdowns of *Dync1i2*, *Dync1li2*, *Dynlrb1* and *Dctn6* reduced the extent of axon outgrowth in cultured DRG neurons (**Figure 1A**), with *Dynlrb1* knockdown providing the most robust effect (**Figure 1A, B**). We then sought to validate this finding in a knockout mouse model for *Dynlrb1* generated by the European Conditional Mouse Mutagenesis Program (EUCOMM) as part of the International Mouse Knockout Consortium. The gene targeted *Dynlrb1^tm1a(EUCOMM)Wtsi^*allele, hereafter referred to as *Dynlrb^tm1a^*, results in a complete knockout of *Dynlrb1* with concomitant expression of β-galactosidase from the same locus; this allele also allows the subsequent generation of a *floxed* conditional allele (*Dynlrb^tm1c^*) upon application of Flp recombinase (**Figure EV1A**)[24]. Homozygous complete knockout mice (*Dynlrb^tm1a/tm1a^*) were not viable, and analyses of early pregnancies showed loss of homozygous null embryos before E9.5, while heterozygous animals were viable. The β-galactosidase expression pattern in heterozygous mice showed that DYNLRB1 was highly expressed in DRGs of E12.5 mouse embryos (**Figure EV1B**). We then crossed (*Dynlrb^tm1a/+^*) heterozygous mice with Thy1-yellow fluorescent protein (YFP) mice (Feng et al., 2000) to allow live imaging of DRG neuron growth in culture. Similarly to *Dynlrb1* knockdown neurons (**Figure 1A, B**), Thy1-YFP-*Dynlrb^tm1a/+^* DRG neurons had significantly reduced axon outgrowth compared to littermate controls (**Figure 1C, D**).

**Figure 1.**
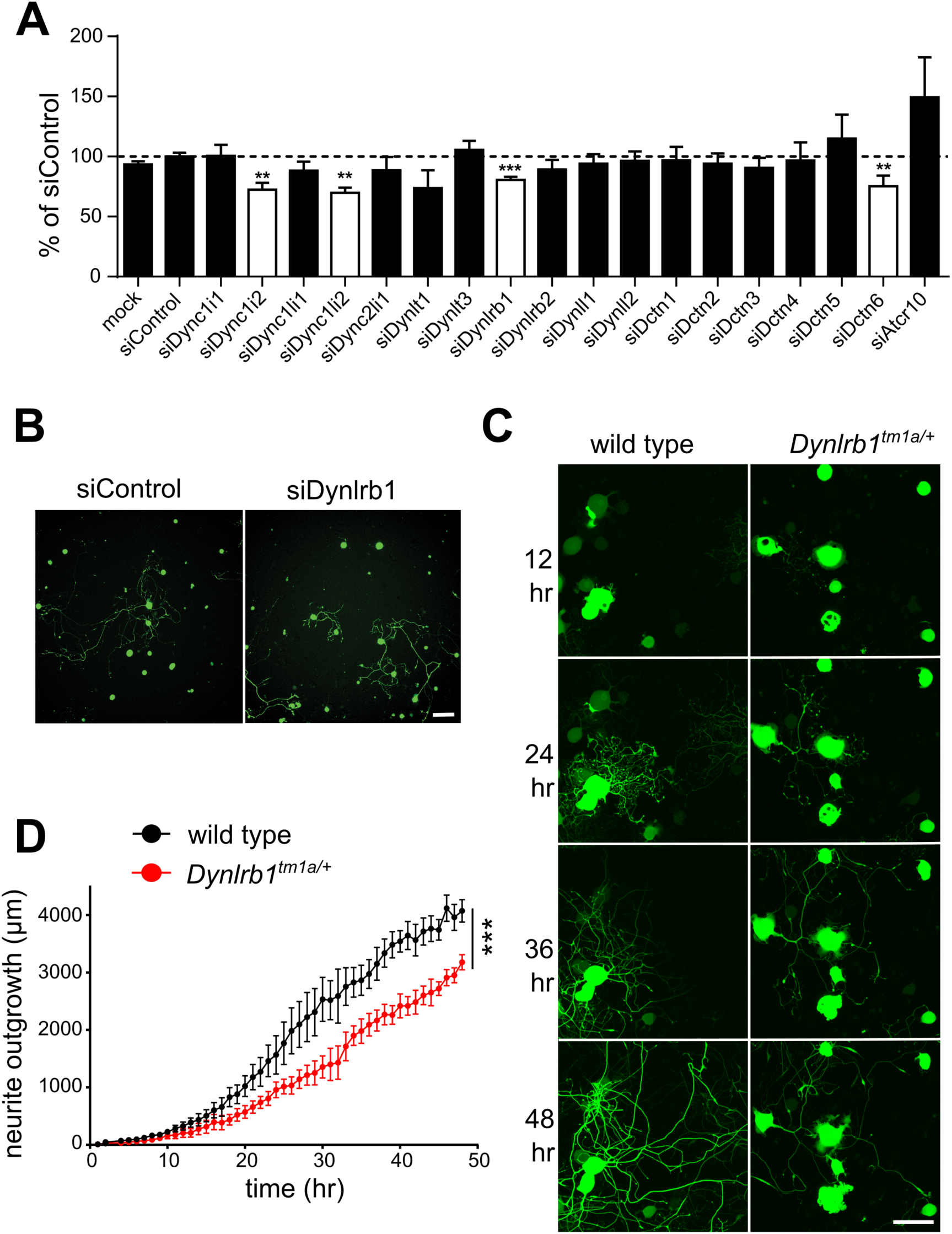
Depletion of DYNLRB1 reduces axon outgrowth in DRG neurons. **A** siRNA screen for effects of dynein complex subunits on neuronal growth (n > 100 cells for each). Positive hits (*Dync1i2, Dync1li2, Dynlrb1 and Dctn6*) were validated in at least three independent experiments (n > 500 cells each). ** p < 0.01, and *** p < 0.001 (Student’s t test and one-way ANOVA). **B** Fluorescence images of cultured DRG neurons from adult Thy1-YFP mice transfected with either siControl or siDynlrb1. Neurons were re-plated 24 hr after siRNA transfection, and images were acquired 48h after re-plating. YFP signal in green. Scale bar, 200 µm. **C** Fluorescent images of cultured DRG neurons from wild type Thy1-YFP mice and Thy1-YFP-*Dynlrb^tm1a/+^* mice (YFP signal in green). Cells were imaged every hour for a period of 48h. Scale bar, 100 µm. **D** Quantification of the time-lapse imaging experiment described in C. Mean ± SEM, *** p < 0.001, n = 3, two-way ANOVA.

**Figure EV1 (related to Figure 1).**
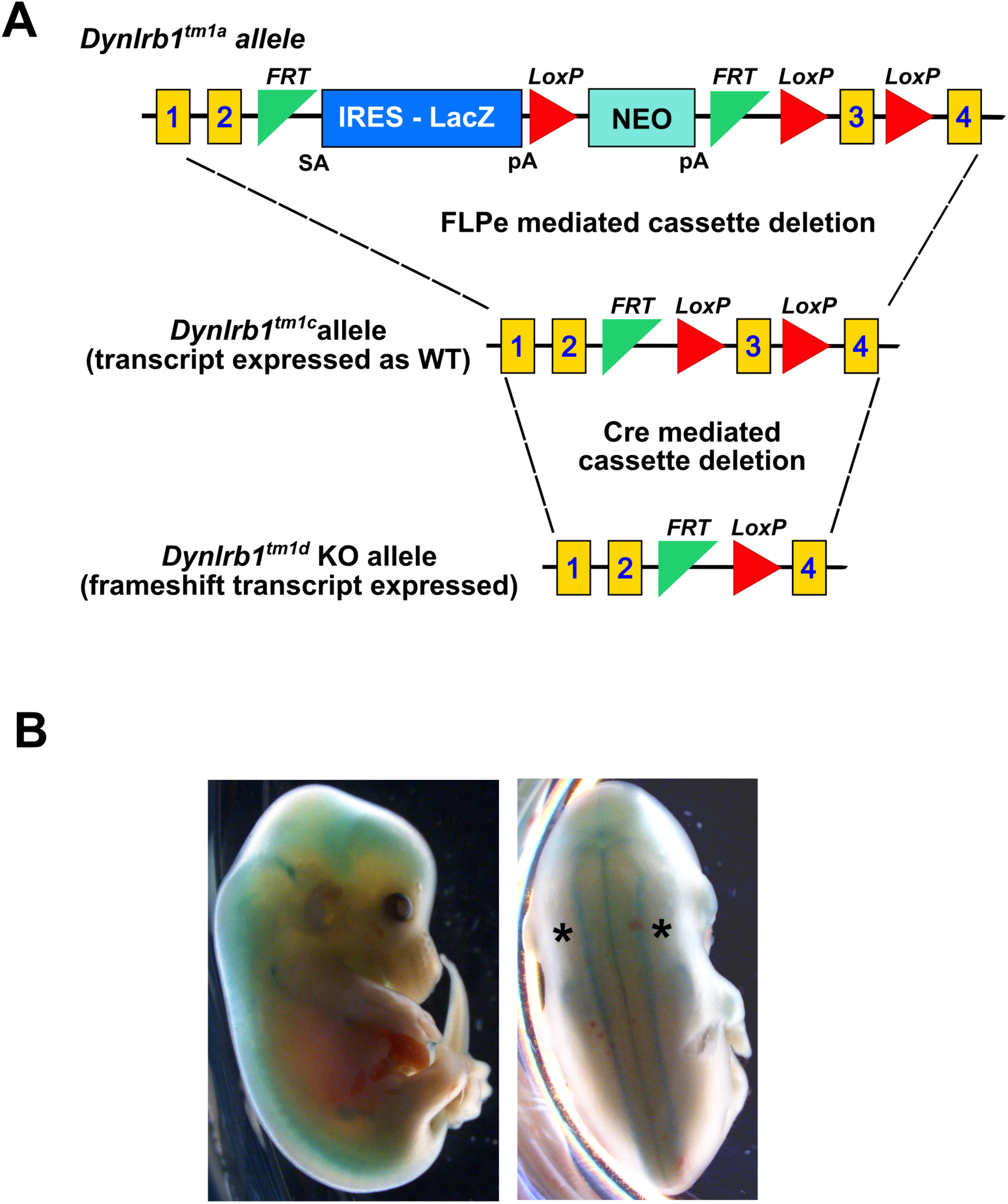
Targeting Vector and Expression Profile of Dynlrb1. **A** Design of the targeting vector of the allele: *Dynlrb1^tm1a(EUCOMM)Wtsi^*from the International Knockout Mouse Consortium (IKMC). The unmodified allele creates a knockout due to the insertion of a β-Gal cassette and a stop codon after exon 2. It can be converted to a floxed ‘Tm1c’ allele by recombining the Frt sites with a *Flp*-driver and to a ‘Tm1d’ KO allele by recombining the LoxP sites with a *Cre*-driver as shown. **B** Lateral and dorsal views of an X-gal-stained E12.5 *Dynlrb^tm1a/+^*embryo to profile expression of β-Gal from the Dynlrb1 locus. Note the strong expression in the nervous system, including the dorsal root ganglia (DRG, pointed at by an asterisk).

### *Dynlrb1* deletion affects survival of proprioceptive neurons

We proceeded to cross *Dynlrb^tm1a/+^*heterozygotes with mice expressing the Flp recombinase [26] to recombine the FRT sites, thereby removing the LacZ sequence [27]. This generated a conditional allele, *Dynlrb^tm1c^* in which *LoxP* sites flank exon 3 of *Dynlrb1* (**Figure EV1A**). *Dynlrb^tm1c^* mice were then crossed with a *RNX3-Cre* driver line [28], to examine the effect of *Dynlrb1* deletion in proprioceptive sensory neurons. In contrast to the full knockout, *RNX3-Dynlrb^tm1d/tm1d^*, hereafter referred to as *RNX3-Dynlrb1*^-/-^, mice were viable to adulthood. However, these mice displayed abnormal hind limb posture (**Figure 2A**), and an uncoordinated walking pattern with abdomen posture close to the ground (**Movie S1**). A battery of tests of motor activity and proprioception showed that *RNX3-Dynlrb1*^-/-^ mice were unable to balance themselves in the rotarod (**Figure 2B**) and exhibited an abnormal walking pattern in the catwalk (**Figures 2C, D**). There was no difference between genotypes in basal motor activity (**Figure EV2A**), but *RNX3-Dynlrb1*^-/-^ mice displayed lower speed and covered less distance than their wild type counterparts in the open field (**Figure EV2B-D**). Overall, these tests reveal a strong impairment of proprioception in adult *RNX3-Dynlrb1*^-/-^ mice.

**Figure 2.**
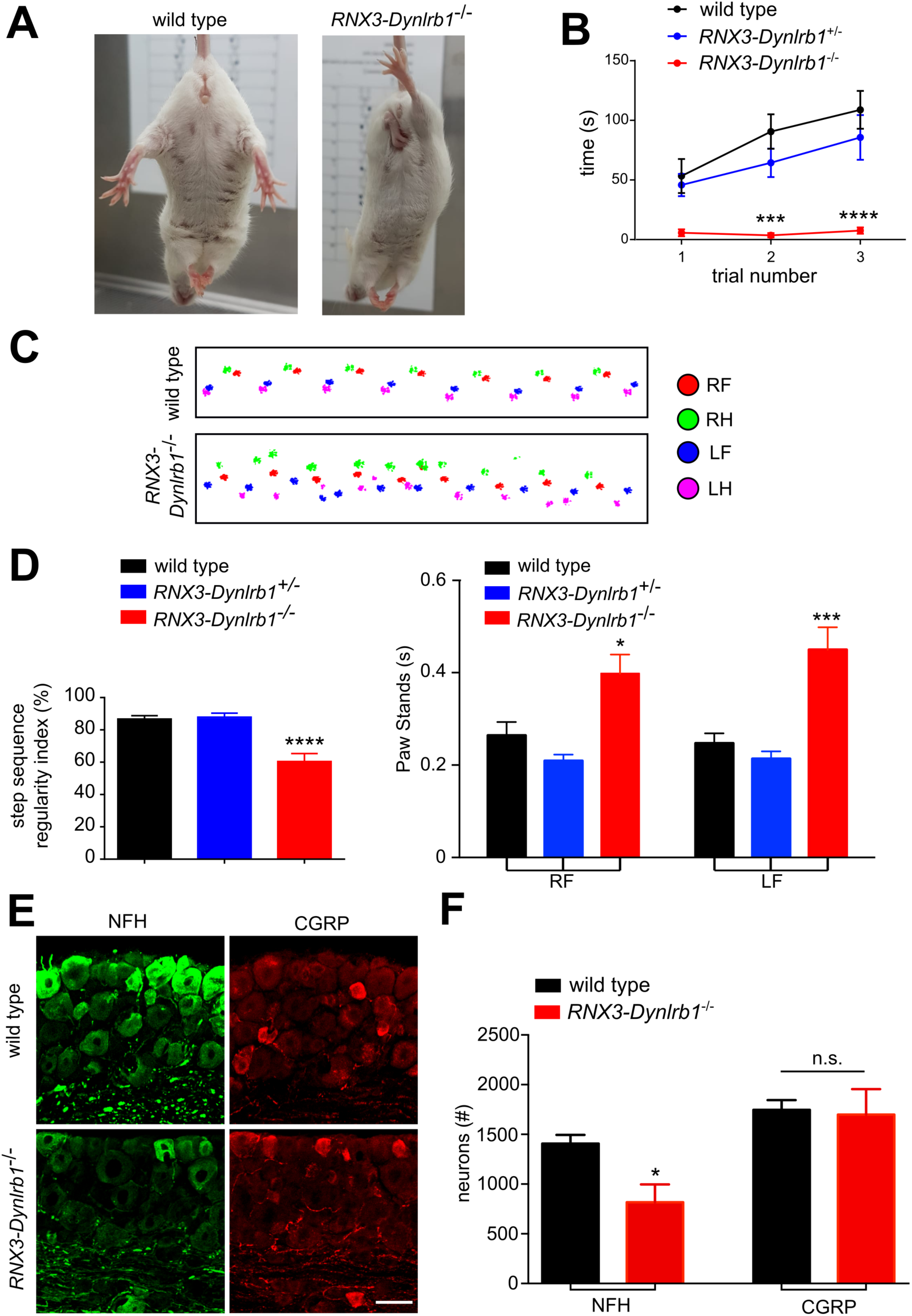
Conditional Dynlrb1 depletion affects survival of proprioceptive neurons in vivo. **A** Abnormal clenched hind-limb phenotype in *RNX-Dynlrb1*^-/-^ mice suspended from the tail, compared to the typical limb extension of wild type animals. **B** Rotarod test (acceleration set at 20 rpm in 240 sec) showing that *RNX-Dynlrb1*^-/-^ mice are unable to balance themselves on the rod. Mean ± SEM, *** p<0.001, n>6 mice per genotype, two-way ANOVA. **C** Representative catwalk gait traces for wild type and *RNX-Dynlrb1*^-/-^ mice. The position of the paws are shown in different colors as indicated, RF - right front, RH - right hind, LF - left front, LH - left hind. **D** Step sequence regularity index and Paw stands from the catwalk gait analyses of C shows impairment in *RNX-Dynlrb1*^-/-^ mice compared to wild type. Mean ± SEM, * p<0.05 *** p<0.001, n>6 mice per genotype, one-way ANOVA followed by Tukey’s HSD post hoc correction for multiple comparisons. **E** Representative DRG ganglia sections from wild type and *RNX-Dynlrb1*^-/-^ mice, stained with either NFH (green) for proprioceptive neurons or CGRP (red) for nociceptors. Scale bar, 50 µm. **F** Quantification of NFH and CGRP positive neurons from the experiment described in G. Mean ± SEM, * p<0.05, n=3 per genotype, unpaired t-test.

**Figure EV2 (related to Figure 2).**
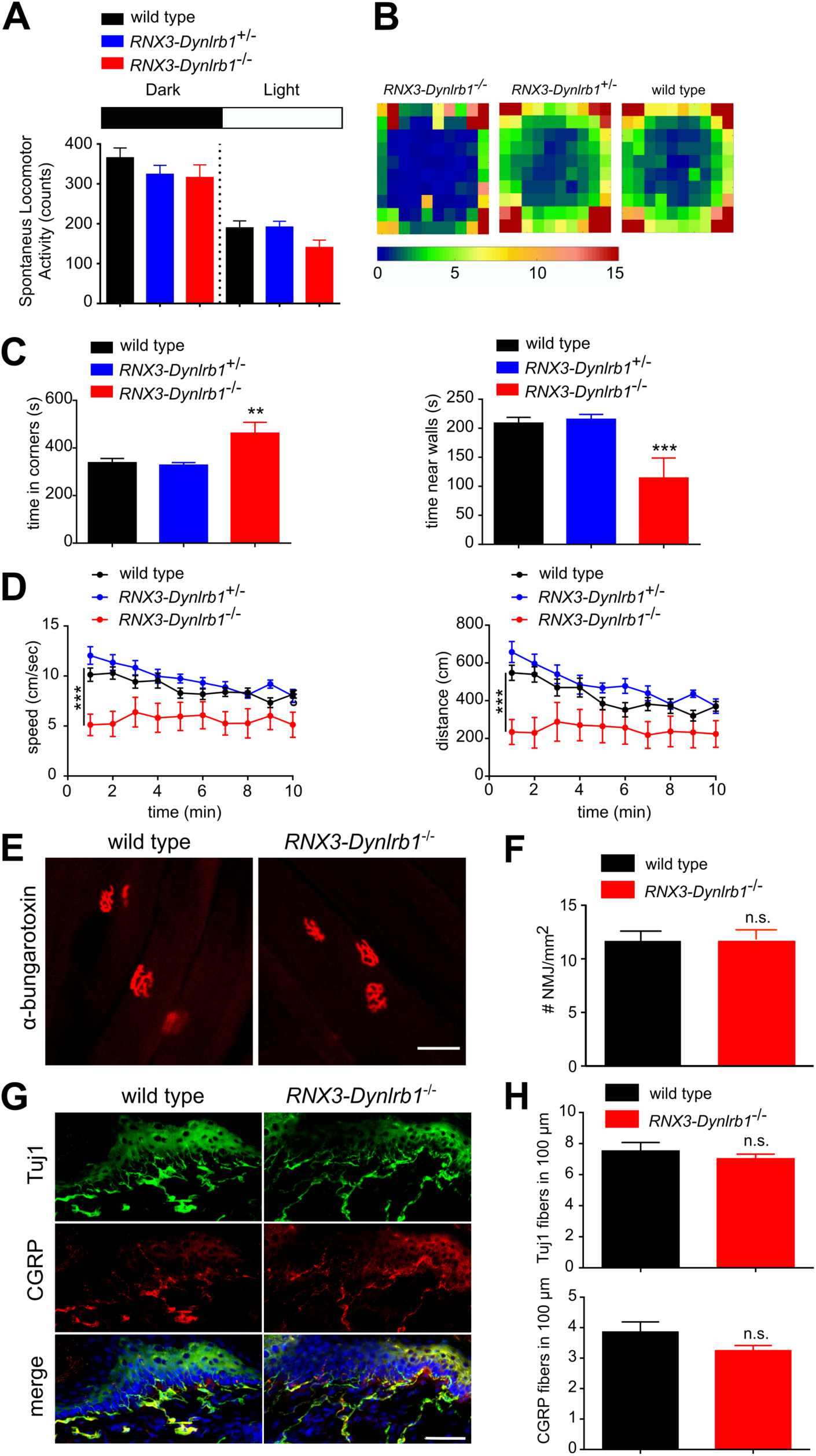
Characterization of RNX3-Dynlrb1 Conditional Knockout Mice. **A** Home cage locomotion analyses show no difference between the genotypes, mean ± SEM, n>6 mice per genotype. **B** Group heatmap representations of the mouse activity (time spent, in seconds) over 10 min of open field exploration of wild type, *RNX3-Dynlrb1*^+/-^ and *RNX3-Dynlrb1*^-/-^ mice. **C** *RNX3-Dynlrb1*^-/-^ mice spend more time in corners and less time near walls in the open-field compared to the other genotypes. Mean ± SEM, ** p<0.01 *** p<0.001, n>6 mice per genotype, one-way ANOVA followed by Tukey’s HSD post hoc correction for multiple comparisons. **D** *RNX3-Dynlrb1*^-/-^ mice also showed reduced speed and distance travelled in the open-field compared to the other genotypes. Mean ± SEM, *** p<0.001, n>6 mice per genotype, two-way ANOVA. **E** α-bungarotoxin staining (red) of the neuromuscular junctions (NMJ) in wild type and *RNX3-Dynlrb1*^-/-^ mice. Scale bar, 50 μm. **F** Quantification of NMJ densities in wild type and *RNX3-Dynlrb1*^-/-^ mice. No significant difference was found between the genotypes, mean ± SEM, n=5 mice per genotype, Unpaired t-test. **G** Sensory innervation of the skin in wild type and *RNX3-Dynlrb1*^-/-^ mice. Tuj1 immunostaining (green) is used to highlight all the nerve endings, while nociceptive neurons are labeled with a CGRP antibody (red). Scale bar, 50 μm . **H** Quantification of Tuj1 and CGRP positive innervation of the skin in wild type and *RNX3-Dynlrb1*^-/-^ mice. No significant difference was found between the genotypes, mean ± SEM, n=4 mice per genotype, unpaired t-test.

To understand the mechanistic basis of this impairment, we performed histological analyses on *RNX3-Dynlrb1*^-/-^ mice. There was no difference between genotypes in neuromuscular junction size or numbers (**Figure EV2E, F**), and in total (Tuj1 positive) or nociceptive (CGRP positive) skin innervation (**Figure EV2G, H**). However, the number of proprioceptive NFH-positive neurons in DRG ganglia was greatly reduced in *RNX3-Dynlrb1*^-/-^ mice (**Figure 2E, F**), while CGRP-positive nociceptive neuron numbers did not differ between genotypes (**Figure 2E, F**). These data reaffirm the specificity of the *RNX3-Cre* driver and indicate that the proprioceptive impairment observed in *RNX3-Dynlrb1*^-/-^ mice is due to a loss of this class of sensory neurons.

### Acute viral-mediated knockdown of *Dynlrb1* causes motor-balance defects *in vivo*

The *RNX3-Cre* driver is activated early in development. In order to determine whether *Dynlrb1* deletion has similar effects upon acute depletion in adult proprioceptive neurons, wild type mice were injected intrathecally with adeno-associated virus (AAV) harboring an shRNA against *Dynlrb1* or an shControl sequence. We used AAV serotype 9 for neuronal-specific knockdown [29, 30], and observed transduction rates of 25-35% in DRG ganglia (**Figure 3A, B**). Efficiency of the knockdown was found to be in the range of 40% decrease in *Dynlrb1* mRNA levels in cultured DRG (**Figure 3C**). 32 days after the injection, mice transduced with AAV-shDynlrb1 developed motor problems and abnormal hind limb posture (**Figure 3D**). AAV-shDynlrb1 transduced mice showed significant impairment in rotarod and catwalk analyses when compared with AAV-shControl transduced animals (**Figure 3E-G**). Overall these data show that *Dynlrb1* is critical for adult sensory neurons and its depletion by knockdown recapitulates the phenotype observed in the *RNX3-Dynlrb1*^-/-^ mice.

**Figure 3.**
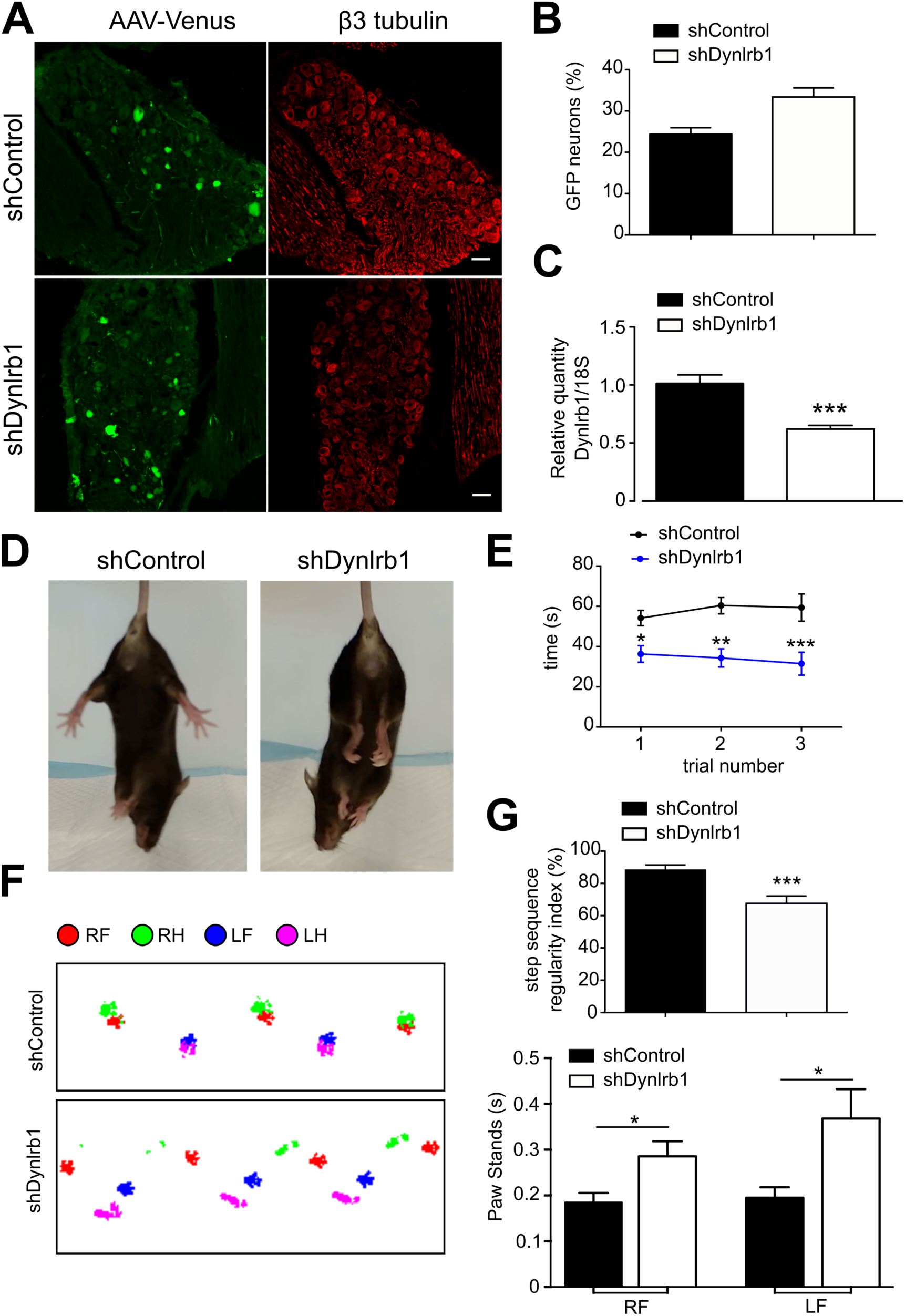
Characterization of Dynlrb1 knockdown in the DRG in vivo. **A** DRG ganglia sections from mice transduced with either AAV9-shControl or AAV9-shDynlrb1. AAV9 infected cells express the Venus reporter (green). Sensory neurons are labeled with β3 tubulin (red). Scale bar, 50 μm. **B** Percentage of GFP-positive neurons normalized to β3 tubulin-positive neurons in DRG sections of the experiment described in A. **C** Quantitative PCRs on RNA extracted from DRG neuron cultures transduced with AAV9-shControl or AAV9-shDynlrb1 for 7 days. AAV9-shDynlrb1 transduced neurons showed 40% reduction of *Dynlrb1* mRNA compared to shControl. Mean ± SEM, *** p<0.001, n=6, unpaired t-test. **D** Tail suspension reveals a clenched hind limb phenotype in mice transduced with AAV9-shDynlrb1 compared to AAV9-shControl-treated animals. **E** Rotarod tests (acceleration set at 20 rpm in 240 sec) on mice transduced as shown, 32 days after viral injection. AAV9-shDynlrb1-transduced mice show significantly reduced performance in the test. Mean ± SEM, * p<0.05, ** p<0.01, *** p<0.001, n>11 mice per group, two-way ANOVA. **F** Representative traces from catwalk gait analyses on mice transduced as described in E. The position of the paws are shown in different colors as indicated, RF – right front, RH - right hind, LF - left front, LH - left hind. **G** Catwalk analysis of experiment described in C. The step sequence regularity index as well as the duration of the paw stands are impaired for both front paws in mice transduced with AAV9-shDynlrb1. Mean ± SEM, * p<0.05, *** p<0.001, n>11 mice per group, unpaired t-test.

### Gene expression changes in *Dynlrb^tm1a/+^* DRG neurons

We then sought to obtain insights on the physiological role(s) of DYNLRB1. Since complete depletion of DYNLRB1 is lethal, we focused on *Dynlrb^tm1a/+^* DRG neurons, which are viable and display reduced neurite outgrowth. RNA-seq of *Dynlrb^tm1a/+^* versus wild type DRG neurons at 12, 24 and 48 hr in culture showed almost no differences between the two genotypes at 12 and 24 hr, while significant differences were observed at 48 hr (**Figure 4A, B, Table S1**). Bioinformatic analyses of the differentially expressed genes dataset revealed changes in a number of signaling and transcriptional pathways, including the canonical Wnt – β-catenin pathway (**Figure 4B, C, Table S2**). Interestingly, direct interaction between a Dynein intermediate chain and β-catenin has been shown in epithelial cells [31], where the complex tethers microtubules at adherens junctions. Thus, DYNLRB1 might be part of a dynein complex carrying β-catenin from the periphery to the nucleus. Indeed, β-catenin nuclear accumulation at 48 h in culture is reduced in *Dynlrb^tm1a/+^*DRG neurons compared to their wild type littermates (**Figure 4D, E**).

**Figure 4.**
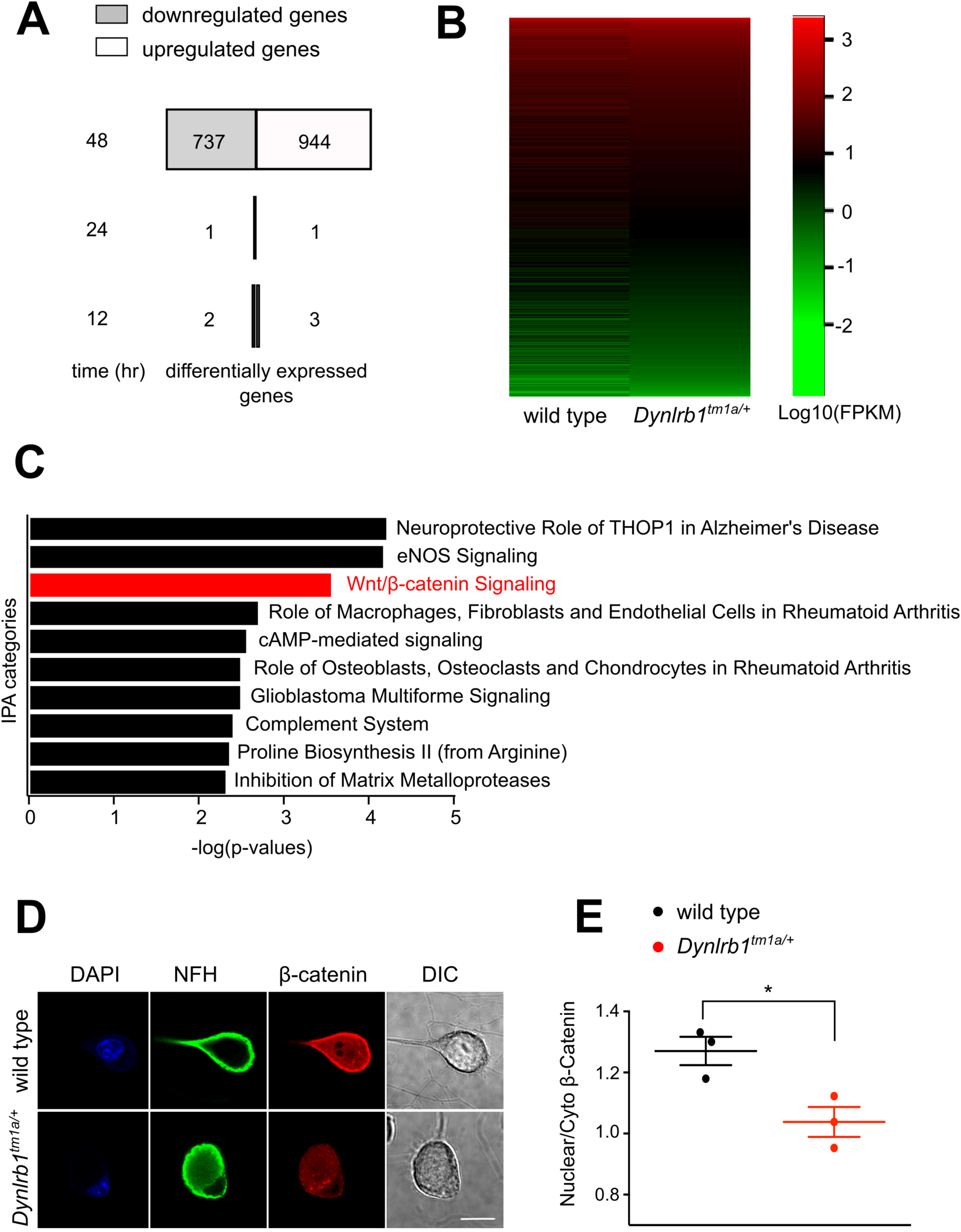
RNA-seq of cultured Dynlrbtm1a/+ DRG neurons highlights signaling deficits. **A** Bar charts representing the numbers of differentially expressed genes (contrast analysis in EdgeR, FDR<0.1) in wild type and *Dynlrb^tm1a/+^*DRG neurons at the indicated time points in culture. **B** Heat map representing the differential expressed genes in wild type and *Dynlrb^tm1a/+^* mice at 48 hr in culture (Log10(FPKM)). **C** Ingenuity pathway analyses on the 48 hr differentially expressed genes dataset, ranked by –log(p values). The Wnt – β−catenin pathway (highlighted in red) was selected for further validation. **D** Representative cell bodies of wild type and *Dynlrb^tm1a/+^*DRG neurons cultured for 48 hr before immunostaining as shown. Scale bar, 20 μm. **E** Quantification of nuclear versus cytoplasmic β−catenin in the experiment described in C. Mean ± SEM, * p<0.05 ** p<0.01, n>3 mice per genotype, one-way ANOVA followed by Tukey’s HSD post hoc correction for multiple comparisons.

### Depletion of *Dynlrb1* has significant effects on axonal retrograde transport

The broad spectrum of gene expression changes in *Dynlrb^tm1a/+^* DRG neurons argues for a general role of this subunit in the dynein complex. We therefore investigated the effect of DYNLRB1 depletion on dynein-mediated axonal retrograde transport, using acute depletion of *Dynlrb1* in culture to circumvent the loss of neurons observed in *RNX3-Dynlrb1*^-/-^ mice. To this end, we transduced *Dynlrb^tm1c/tm1c^*DRG neurons with Adenovirus 5 (Ad5) expressing a Cre-GFP, to induce recombination of the *Dynlrb^tm1c^* allele to *Dynlrb^tm1d^*in infected cells. We first optimized the transduction protocol using neurons from the Ai9 mouse [32], which expresses td-Tomato under the control of a LoxP-flanked STOP cassette and showed that 72 hr of incubation with Ad5 Cre-GFP are sufficient to express the Cre and trigger the recombination of LoxP sites in cultured DRG neurons (**Figure EV3A, B**). We then tracked axonal retrograde transport of acidic organelles, including lysosomes, late endosomes and autophagosomes in Ad5-Cre: *Dynlrb^tm1d/tm1d^*DRG neurons using Lysotracker Red. Indeed, Ad5-Cre: *Dynlrb^tm1d/tm1d^*transduced neurons 96 h in culture showed a significant impairment of axonal retrograde transport of these organelles, both in terms of velocity of individual carriers and the overall fraction of moving carriers (**Figure 5A-C**).

**Figure 5.**
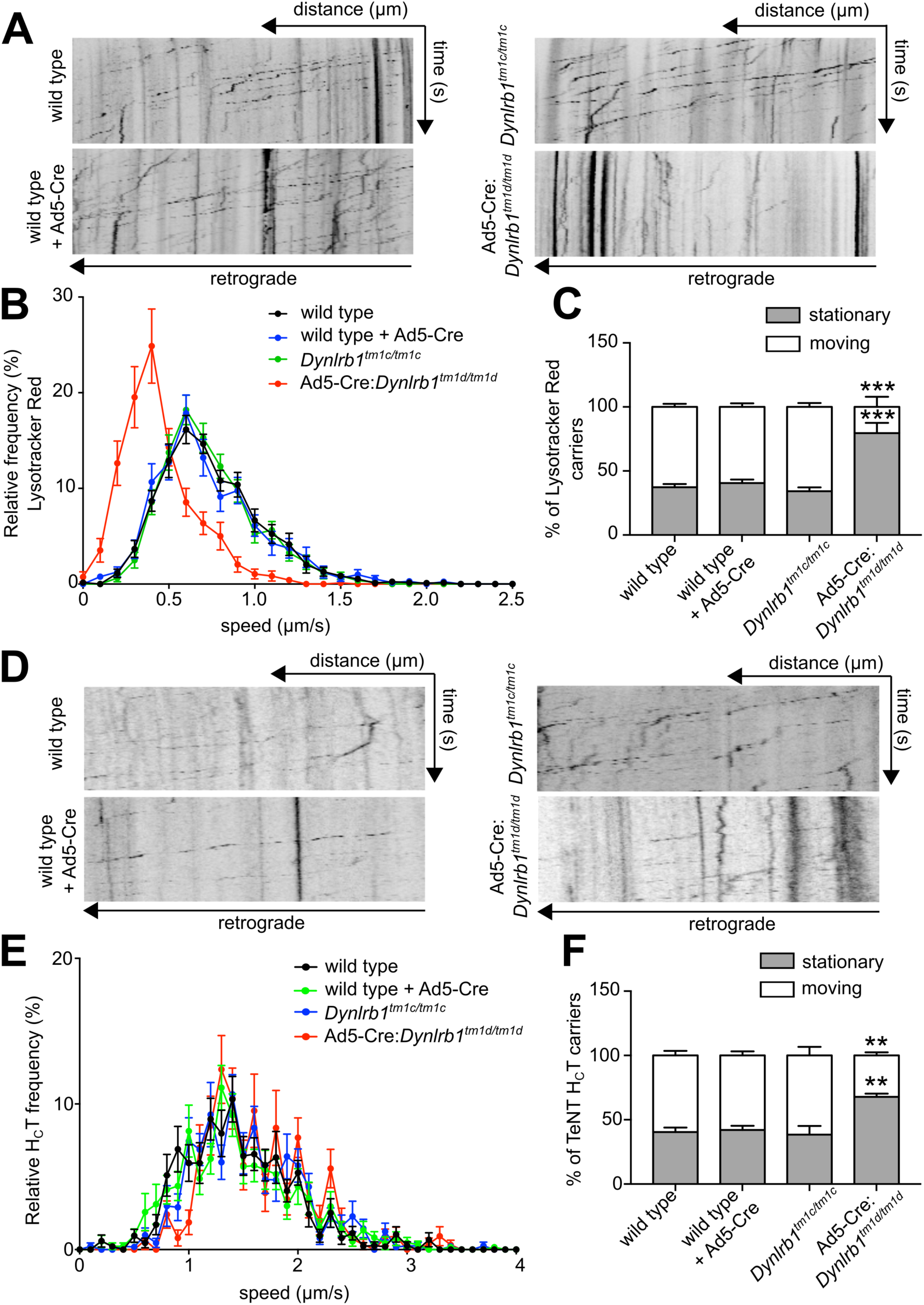
Axonal retrograde transport deficits after Dynlrb1 depletion. **A** Representative kymographs of Lysotracker Red tracking on wild type and *Dynlrb^tm1c/tm1c^*DRG neurons transduced for 96h in culture as indicated (Ad5-Cre, Adenovirus5 expressing Cre-GFP) for 96 hr. Diagonal lines indicate retrogradely moving carriers. **B** Velocity distributions from the experiment shown in A, Mean ± SEM, n>32 axons per group over 3 independent biological repeats. **C** Fraction of moving versus stationary carriers in the experiment described in A. Mean ± SEM, *** p<0.001, n>32 axons per group over 3 independent biological repeats, two-way ANOVA. **D** Representative kymographs of TeNT H_C_T tracking on wild type and *Dynlrb^tm1c/tm1c^* DRG neurons transduced for 96 hr in culture as indicated. **E** Velocity distributions from the experiment shown in D. Mean ± SEM, n>36 axons per group over 3 independent biological repeats. **F** Fraction of moving versus stationary carriers in the experiment described in D. Mean ± SEM, ** p<0.01, n>36 axons per group over 3 independent biological repeats, two-way ANOVA.

**Figure EV3 (related to Figure 4).**
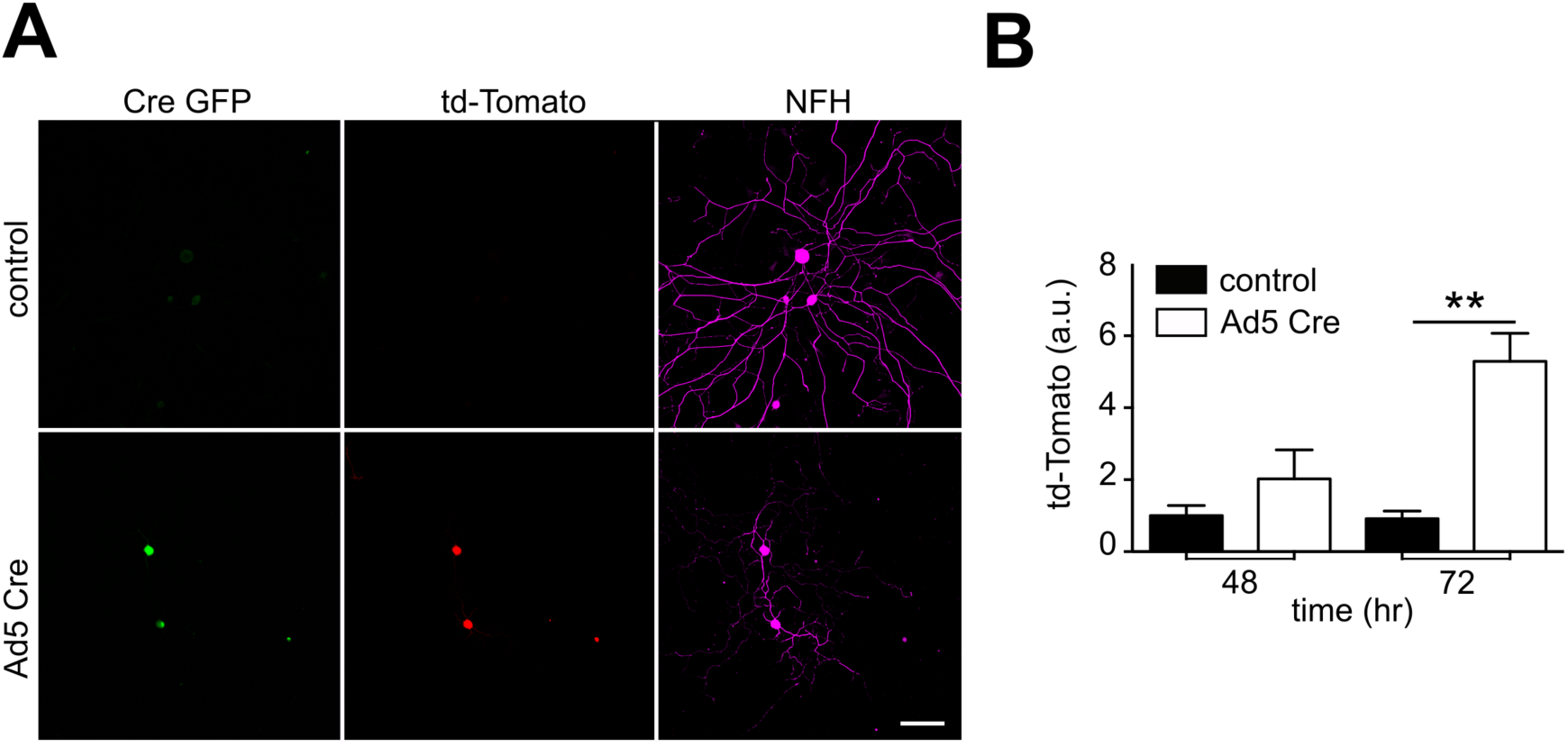
Transduction Efficiency with Adenovirus-Cre. **A** DRG neurons from Ai9 mice 72 hr after transduction with Cre-GFP Adenovirus. Transduced cells are positive for the antibody α-GFP (green) and td-Tomato (red). Neurons are identified with the cytoskeletal marker NFH (magenta). Scale bar, 100 μm. B Quantification of relative td-Tomato expression in the experiment described in A. Mean ± SEM, ** p<0.01, n = 3, one-way ANOVA followed by Tukey’s HSD post hoc correction for multiple comparisons.

We further examined transport properties using a marker for neurotrophin signaling endosomes, the binding fragment of tetanus neurotoxin (H_C_T) fragment [33, 34]. Whilst we could not detect any significant differences in the rates of transport of H_C_T-positive retrograde axonal carriers in Ad5-Cre: *Dynlrb^tm1d/tm1d^* neurons (**Figure 5D, E**), the percentage of stationary carriers was significantly increased (**Figure 5F**), as was also found for Lysotracker Red-positive organelles (**Figure 5C**). Thus, depletion of DynlRb1 has significant effects on retrograde transport of a broad spectrum of dynein cargos.

## DISCUSSION

The dynein complex has essential roles in all multicellular organisms, most prominently in long distance axonal transport in neurons. Studies on specific roles of dynein carriers have suffered from a lack of appropriate mouse models, since homozygous deletion or mutation of *Dync1h1* results in early embryonic death (Hafezparast et al., 2003; Harada et al., 1998). We have characterized a new conditional knockout model for the DYNLRB1 subunit of dynein and use this model to show that DYNLRB1 is required for retrograde axonal transport of a broad spectrum of cargos in sensory axons, as well as for neuronal survival. These findings were surprising given the notion that DYNRLB1 should function mainly as a binding adaptor for TGFβ signaling complexes and other specific cargos [11,14–18], whereas our data suggests that DYNLRB1 is essential for dynein-mediated axonal transport of lysosomes, late endosomes and signaling endosomes.

The hindlimb proprioception phenotype observed in both *RNX3-Dynlrb1*^-/-^ mice and *Dynlrb1* viral-mediated knockdown is highly reminiscent of dynein heavy chain mutant models, such as *Dync1h1^Loa^*and *Dync1h1^Cra1^* [35, 37]. Both models show deficits in endosomal axonal retrograde transport [35] and proprioception [37, 38], as we have shown above for knockout or knockdown of *Dynlrb1*. In addition, depletion of *Roadblock* in *Drosophila* was described to cause defects in intra-axonal accumulation of cargoes and severe axonal degeneration [12]. The distal enrichment of axonal accumulations observed in *Roadblock* mutants was suggested to arise from axonal transport deficits originating at the synapse [12]. Moreover, incorporation of DYNLRB1 and DYNLRB2 into the Dynein-2 complex, which drives retrograde protein trafficking in primary cilia, was recently shown to be crucial for dynein-2 transport function [39]. While the precise mechanism by which dynein activity is impaired upon loss of DYNLRB1 is yet to be determined, it is interesting to note that this subunit can bind directly to core components of the dynein complex such as the intermediate chains DYNCI1 and DYNCI2 (IC74-1 and IC74-2) [13]. The latter directly associate with the dynactin complex, which facilitates interactions of numerous cargos with the main motor complex, and expedites processive transport along microtubules (Karki and Holzbaur, 1995; Reck-Peterson et al., 2018).

Our findings show that *Dynlrb1* is critical not only for embryonic development, but also for maintenance of sensory neurons in adulthood. Deficits in dynein functions have been implicated in the pathogenesis of neurodegenerative diseases such as Amyotrophic Lateral Sclerosis (ALS) [40–45], and an increasing number of human neurodegenerative pathologies have been linked to mutations of dynein complex genes [46]. In addition, mutations and/or disruption of the function of the dynactin complex has been shown to be connected to motor neuron degeneration and ALS [47–51]. The new mouse model described herein will expand the options for studying such roles of dynein, using conditional targeting of Dynlrb1 for both temporal and tissue/cell type specificity in such studies *in vivo*.

## Acknowledgments

We thank Nicolas Panayotis for expert guidance and advice with the behavioral assays; Nitzan Korem and Nataliya Okladnikov for excellent technical assistance; Vladmir Kiss for professional microscopy support; Dalia Gordon, Indrek Koppel, Michael Tsoory and Stefanie Alber for helpful comments and advice; Avraham Yaron for the kind gift of the Ai9 mice and Avri Ben-Ze’ev for the kind gift of a β-catenin antibody. We thank the Medical Research Council Mammalian Genetics Unit, Harwell, UK for providing the *Dynlrb1^tm1a(EUCOMM)Wtsi^* mice.

## Author contributions

M.F. and M.T. designed the study. M.T., A.D.P., I.R., L.M., P.D.M., and R.K. performed experiments and data analyses. R.K. carried out RNA sequencing. M.F., E.M.C.F., G.C., and G.S. supervised research. M.F. and M.T. wrote the initial manuscript draft. All authors revised the manuscript and approved the final version.

## SUPPLEMENTARY MOVIE, TABLES AND THEIR LEGENDS

**Source movie 1, related to Figure 1.** Video of a 2 months old *RNX3-Dynlrb1*^-/-^ mouse showing the hindlimb clenching as well as the abnormal walk and posture (abdomen close to the ground).

**Supplementary Table 1, related to Figure 3.** List of differentially regulated genes highlighted by RNA sequencing between wild type and *Dynlrb1*^+/-^ mice at 48h. The genes are ranked by FDR. An additional sheet is present, with the list of genes selected by a value of FDR<0.1. Raw and processed data were deposited within the Gene Expression Omnibus (GEO) repository (www.ncbi.nlm.nih.gov/geo), accession number (GSE131455).

**Supplementary Table 2, related to Figure 3.** List of pathways identified by the Ingenuity Pathway Analysis (QIAGEN Bioinformatics, sheet1). A list of all the genes of the WNT pathway found to be differentially regulated and the value of the change fold is also shown (sheet2).

## MATERIAL AND METHODS

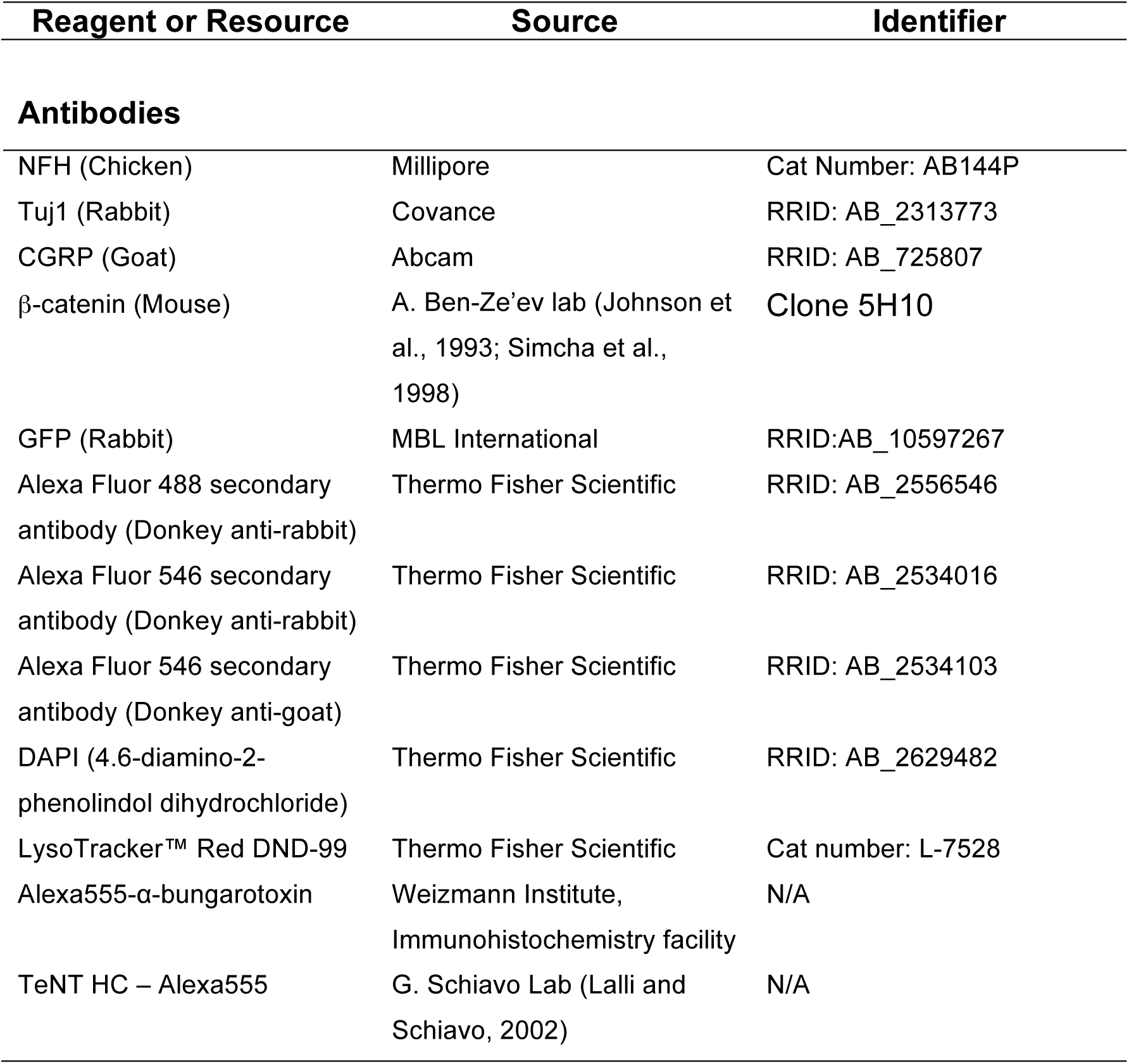
Key Resource Table

Key Resource Table
| Reagent or Resource | Source | Identifier |
| --- | --- | --- |
| **Antibodies** |  |  |
| NFH (Chicken) | Millipore | Cat Number: AB144P |
| Tuj1 (Rabbit) | Covance | RRID: AB_2313773 |
| CGRP (Goat) | Abcam | RRID: AB_725807 |
| β-catenin (Mouse) | A. Ben-Ze'ev lab [52,53] | Clone 5H10 |
| GFP (Rabbit) | MBL International | RRID:AB_10597267 |
| Alexa Fluor 488 secondary antibody (Donkey anti-rabbit) | Thermo Fisher Scientific | RRID: AB_2556546 |
| Alexa Fluor 546 secondary antibody (Donkey anti-rabbit) | Thermo Fisher Scientific | RRID: AB_2534016 |
| Alexa Fluor 546 secondary antibody (Donkey anti-goat) | Thermo Fisher Scientific | RRID: AB_2534103 |
| DAPI (4.6-diamino-2-phenolindol dihydrochloride) | Thermo Fisher Scientific | RRID: AB_2629482 |
| LysoTracker™ Red DND-99 | Thermo Fisher Scientific | Cat number: L-7528 |
| Alexa555-α-bungarotoxin | Weizmann Institute, Immunohistochemistry facility | N/A |
| TeNT HC – Alexa555 | G. Schiavo Lab [34] | N/A |

Experimental Models: Organism/Strains

Experimental Models: Organism/Strains
|  |  |  |
| --- | --- | --- |
| Mouse/C57BL/6OlaHsd | Harlan Israel | N/A |
| Mouse/C57BL/6OlaHsd/*Dynlrb1*<sup>tm1a</sup> | International Knockout Mouse Consortium | allele: *Dynlrb1*<sup>tm1a(EUCOMM)Wtsi</sup> |
| Mouse/FLP1eR - 129 / 129S4/SvJaeSor-Gt(ROSA)26Sor<sup>tm1(FLP1)Dym/J</sup> | [26] | N/A |
| Mouse/Ai9 | [32] | N/A |
| Mouse/R26LacZ/Runx3<sup>Cre</sup> | [28] | N/A |

siRNA screen Dharmacon smartpools

siRNA screen Dharmacon smartpools
| Catalog Number | Gene Symbol | Locus ID | Accession |
| --- | --- | --- | --- |
| D-001210-02 | Non-targeting siRNA | N/A | N/A |
| M-043143-01 | *Dync1i1* | 13426 | NM_010063 |
| M-043171-01 | *Dync1i2* | 13427 | NM_010064 |
| M-051314-01 | *Dync1li1* | 235661 | NM_146229 |
| M-061580-01 | *Dync1li2* | 234663 | NM_001013380 |
| M-062961-01 | *Dync2li1* | 213575 | NM_172256 |
| M-055779-01 | *Dynlt1* | 21648 | NM_009342 |
| M-059901-01 | *Dynlt3* | 67117 | NM_025975 |
| M-044820-01 | *Dynlrb1* | 67068 | NM_025947 |
| M-050822-01 | *Dynlrb2* | 75465 | NM_029297 |
| M-063157-01 | *Dynll1* | 56455 | NM_019682 |
| M-059592-01 | *Dynll2* | 68097 | NM_026556 |
| M-044821-00 | *DCTN1* | 13191 | NM_007835 |
| M-056851-00 | *DCTN2* | 69654 | NM_027151 |
| M-063002-00 | *DCTN3* | 53598 | NM_016890 |
| M-063074-00 | *DCTN4* | 67665 | NM_026302 |
| M-063270-00 | *DCTN5* | 59288 | NM_021608 |
| M-047876-00 | *DCTN6* | 22428 | NM_011722 |
| M-049777-00 | *ACTR10* | 56444 | NM_019785 |

### Contact for Reagent and Resources Sharing

Further information and requests for resources and reagents should be directed to Marco Terenzio.

### Experimental Model and Subject Details

#### Animal Subjects

All animal procedures were approved by the IACUC of the Weizmann Institute of Science. The knockout mouse model for *Dynlrb1* was generated by the European Conditional Mouse Mutagenesis Program (EUCOMM) as part of the International Mouse Knockout Consortium (IKMC, allele: Dynlrb1^tm1a(EUCOMM)Wtsi^) using a KO first targeting vector (MPGS8001_A_B08, IKMC Project 25156) [24]. All mouse strains used were bred and maintained at the Veterinary Resource Department of the Weizmann Institute of Science. For live imaging of DRG outgrowth (**Figure 1A-D**) C57BL/6J YFP16 mouse were used [25]. All control mice were age-, sex-matched littermates. For testing Adenovirus5 Cre-GFP infectivity Ai9 mice were used [32]. Mice utilized for behavioral experiments were kept at 24.0 ± 0.5°C in a humidity-controlled room under a 12-h light– dark cycle with free access to food and water. Experiments were carried out on 2-6 months old male mice, unless specified otherwise.

### Generation of the *Dynlrb^tm1c^* and RNX3-*Dynlrb^tm1d/tm1d^* lines

The *Dynlrb1* mice acquired from the International Mouse Knockout Consortium (IKMC, allele: Dynlrb1^tm1a(EUCOMM)Wtsi^).The gene targeted *Dynlrb1^tm1a(EUCOMM)Wtsi^*allele, hereafter referred to as *Dynlrb^tm1a^*, results in a complete knockout of *Dynlrb1* with concomitant expression of β-galactosidase from the same locus; this allele also allows the subsequent generation of a *floxed* conditional allele (*Dynlrb^tm1c^*) upon application of the *Flp* recombinase [24]. *Dynlrb^tm1a/+^* heterozygous mice were crossed with a mouse harboring the *Flp* recombinase in all tissues (FLP1eR - 129/129S4/SvJaeSor-Gt(ROSA)26Sor^tm1(FLP1)Dym^/J) [26], which was provided by the veterinary services of the Weizmann Institute of Science to recombine the FRT sites and so remove the LacZ sequence and giving a conditional allele, *Dynlrb^tm1c^* in which *LoxP* sites flank a likely critical exon, exon 3, of *Dynlrb1* ([24], Supplementary Figure 1A). We further crossed the *Dynlrb^tm1c^* mice with the proprioceptive Cre-driver R26LacZ/Runx3^Cre^ [28] to generate the *RNX3-Dynlrb^tm1d/tm1d^*line, which we refer to in the text as *RNX3-Dynlrb1* (Supplementary Figure 1A) [24], in which *Dynlrb1* is depleted in proprioceptive neurons.

## Methods

### DRG Culture

Adult mice DRGs were dissociated with 100 U of papain followed by 1 mg/ml collagenase-II and 1.2 mg/ml dispase. The ganglia were then triturated in HBSS, 10 mM Glucose, and 5 mM HEPES (pH 7.35). Neurons were recovered through Percoll, plated on laminin, and grown in F12 medium (ThermoFisher Scientific) as previously described [54–56].

### siRNA Screen

Adult C57BL/6 YFP16 mouse DRG [25] were dissociated and an siRNA screen was conducted as previously described [23]. Briefly, mouse siGenome smart pools (Dharmacon) were used for transfection of neuronal cultures two hours after plating, using DharmaFect 4 as the transfection reagent. 24 hours after siRNA transfection the neurons were plated in new poly-L-lysine and Laminin treated glass bottom 96 well plates at the same density (80,000-100,000 cells per well) and imaged 72 h after transfection. Images were captured at X10 magnification on ImageXpress Micro (Molecular Devices) and analyzed using the Metamorph software (Molecular Devices). The total outgrowth parameter was considered, defined as the sum of lengths of all processes and branches per cell. Statistical analyses were carried out with Student’s t-test and One Way ANOVA using the statistical software GraphPad Prism.

### Growth rate analysis

Adult C57BL/6J YFP16 mouse DRG [25] were dissociated as previously described and neurons were plated on MatTek glass bottom dishes (MatTek corporation). YFP6 - Wild type and YFP16-*Dynlrb1*^+/-^ neurons were imaged at 1h intervals in a Fluoview (FV10i, Olympus) automated confocal laser-scanning microscope with built-in incubator chamber. Neuronal morphology was subsequently quantified using the Metamorph sofware (Molecular Devices). The total outgrowth parameter was considered, defined as the sum of lengths of all processes and branches per cell. Statistical analyses were carried out with a Two-way ANOVA test using the statistical software GraphPad Prism.

### Adenovirus infection of DRG neurons from Ai9 mice

The efficiency transduction of adenovirus 5 Cre-GFP (Ad5 Cre-GFP) and the efficiency of the viral Cre were tested by infecting DRG neurons with Ad5 Cre-GFP at a multiplicity of infection (MOI) of 100. DRG neurons from Ai9 mice [32] were dissected as previously described. The Ai9 Cre reporter mice harbour a LoxP-flanked STOP cassette preventing transcription of a CAG promoter-driven red fluorescent protein variant (td-Tomato). The virus was added directly into the F12 culture medium immediately after plating. Infected cultures were then incubated at 37°C for 48 and 72 h. After each time-point, neurons were washed in PBS, fixed in 4% PFA in PBS for 30 min and immunostained with an α-GFP antibody (1:1000) as previously described. Coverslips were then mounted with Flouromount-G^TM^ and imaged using the ImageXpress (Molecular Devices) automated microscope at 10X magnification or using a confocal laser-scanning microscope (Olympus FV1000, 60x oil-immersion objective Olympus UPLSAPO - NA 1.35). Image quantification was performed using the proprietary software Metamorph (Molecular Devices). Statistical analysis was carried out with a one-way ANOVA followed by Tukey post-hoc multiple comparison test using the statistical software GraphPad Prism.

### Axonal transport experiments

Wild type and *Dynlrb1 Flox*^-/-^ DRG ganglia were dissociated as previously described and neurons were plated on MatTek glass bottom dishes (MatTek corporation) in F12 medium (ThermoFisher Scientific) supplemented with 10 μM cytosine β-D-arabinofuranoside hydrochloride (AraC, Sigma-Aldrich, C6645) to inhibit glial proliferation. Adenovirus5 Cre-GFP was added to the neurons 4 h after plating. 96 h after infection with the virus, cells were washed in warm HBSS medium (ThermoFisher Scientific). LysoTracker™ Red DND-99 (ThermoFisher Scientific) (200 nM concentration) or AlexaFluor555-conjugated H_C_T (40 nM) were added to the cells. Neurons were transferred to a a Fluoview (FV10i, Olympus) automated confocal laser-scanning microscope with built-in incubator chamber and imaged with a 60X water immersion objective. 120 frames at intervals of 5s between each frame were taken for every transport movie. Movies were tracked manually using the Manual Tracking plugin of the ImageJ software; while the percentage of stationary vs moving carriers was performed on kymographs generated using the Multi Kymograph plugin of ImageJ. Statistical analysis was performed with two-way ANOVA using the statistical software GraphPad Prism.

### β-Catenin nuclear accumulation in *Dynlrb1*^+/-^ DRG neurons

Wild type and *Dynlrb1^+/-^* DRG ganglia were dissociated as previously described and cells were plated on coverslips in F12 medium (ThermoFisher Scientific). 48 h after plating cells were washed in PBS, fixed in 4% PFA in PBS for 30 min and immunostained for β−catenin (1:100 dilution), NFH (1:2,000 dilution) and DAPI as described previously. Coverslips were then mounted with Flouromount-G^TM^ and imaged using a confocal laser-scanning microscope (Olympus FV1000, 60x oil-immersion objective Olympus UPLSAPO - NA 1.35). Image quantification was performed using ImageJ software. Nuclei were identified using a mask drawn on the DAPI straining and cytoplasm was masked using the NFH staining. Nuclear and cytoplasmic intensity of β−catenin was recorded. Statistical analysis was carried out with a one-way ANOVA followed by Tukey post-hoc multiple comparison test using the statistical software GraphPad Prism.

### Design and production of AAV shDynlrb1

Design of AAV shRNA constructs was based on AAV-shRNA-ctrl (addgene, plasmid #85741, [57]). Sh-Ctrl sequence was replaced by a sequence targeting *Dynlrb1* using the BamHI and XbaI restriction sites. The primers used for cloning were:

Forward: GATCCGATCCAGAATCCAACTGAATAGAAGCTTGTATTCAGTTGGATTCTGGATCTT TTTTT,

Reverse: CTAGAAAAAAAGATCCAGAATCCAACTGAATACAAGCTTCTATTCAGTTGGATTCTG GATCG

Purified adeno associated virus (AAV) was produced in HEK 293t cells (ATCC®), with the AAVpro® Purification Kit (All Serotypes) from TaKaRa (#6666). For each construct 10 plates (15 cm) were transfected with 20 µg of DNA (AAV-plasmid containing our construct of interest + 2 helper plasmids for AAV9) using jetPEI® (Polyplus) in DMEM medium without serum or antibiotics. Medium (DMEM, 20 % FBS, 1 mM sodium pyruvate, penicillin-streptomycin) was added on the following day to a final concentration of 10% FBS and extraction was done 3 days post transfection. Purification was performed according to the manufacturers’ instructions (TaKaRa, #6666).

### Immunohistochemistry of tissue samples

Tissue samples from muscle, skin, DRGs and sciatic nerve were harvested and fixed in 4% PFA in PBS (30 min for skin at RT, overnight at 4°C for muscles, DRG ganglia and sciatic nerve). The tissues were then washed in PBS and equilibrated in 30% w/v sucrose in PBS, embedded in Optimal Cutting Temperature Compound (O.C.T., Tissue-Tek®) and finally cryo-sectioned at a thickness ranging from 10-50 μm. Sections were then re-hydrated in PBS, blocked and permeabilized with 5% goat serum, 1% BSA, 0.1% Triton X-100 in PBS, or donkey serum, 0.1% Triton X-100 in PBS for 1 h and incubated with primary antibody overnight at 4°C. Sections were then washed 3 times in PBS and incubated for 2 h with secondary antibodies from Jackson Immunoresearch (Alexa Fluor 647 or Alexa Fluor 488 conjugated, 1:500 dilution in PBS). Slides were then washed in PBS, mounted with Flouromount-G^TM^, and imaged using a confocal laser-scanning microscope (Olympus FV1000, 60x oil-immersion objective Olympus UPLSAPO - NA 1.35).

### NMJ plate counting

Gastrocnemius muscles from wild type and *RNX3-Dynlrb1*^-/-^ mice were sectioned at a thickness of 50 μm. Staining of the neuromuscular junction was achieved as previously described by incubation with Alexa 555-α-bungarotoxin (1:1000 dilution). Sections were then imaged using a confocal laser-scanning microscope (Olympus FV1000, 60x oil-immersion objective Olympus UPLSAPO - NA 1.35) and the number of positive NMJ-plate per area were manually counted and normalized by the area. Statistical analysis was performed with t-test using the statistical software GraphPad Prism.

### Sensory skin innervation

Hind paw skin of wild type and *RNX3-Dynlrb1*^-/-^ mice was sectioned at a thickness of 20 μm and immunostained as previously described with an α-CGRP and α- TUJ1 antibodies. The nuclear counterstaining DAPI was added to highlight all cells in the tissue. Sections were imaged using a confocal laser-scanning microscope (Olympus FV1000, 60x oil-immersion objective Olympus UPLSAPO - NA 1.35). The number of nerve endings was manually counted and normalized by the area of the skin. Statistical analysis was performed with t-test using the statistical software GraphPad Prism.

### DRG neuron count

L4 DRGs from wild type and *RNX3-Dynlrb1*^-/-^ mice were serially sectioned at a thickness of 20 μm in set of 3. One set for each DRG was then processed for immunostaining as previously described for α-NFH, α-CGRP and α-TUJ1 antibody [58]. NFH and CGRP positive neurons were manually counted in blind. Statistical analysis was performed with t-test using the statistical software GraphPad Prism.

### Gene expression analysis by Q-PCR

Wild type DRG ganglia were dissociated as described above and neurons were plated in F12 medium (ThermoFisher Scientific) supplemented with 10 μM cytosine β-D-arabinofuranoside hydrochloride (AraC, Sigma-Aldrich, C6645) to inhibit glial proliferation. AAV9-shControl and AAV9-shDynlrb1 viruses were added to the neurons 4 h after plating. 7 days after infection, cells were washed in PBS and total RNA was extracted using the Oligotex mRNA Mini Kit (Qiagen). RNA purity and concentration was determined, and 300 ng of total RNA was then used to synthesize cDNA using SuperScript III (Invitrogen). Q-PCR was performed on a ViiA7 System (Applied Biosystems) using PerfeCTa SYBR Green (Quanta Biosciences, Gaithersburg, USA). Forward/Reverse primers were designed to span exon-exon junction, and the RNA was treated with DNase H to avoid false-positives. Amplicon specificity was verified by melting curve analysis. All Q-PCR reactions were conducted in technical triplicates and the results were averaged for each sample, normalized to 18S levels and analyzed using the comparative ΔΔCt method (Livak and Schmittgen, 2001). The primers (*Mus musculus*) that were used are the following:

18S forward: AAACGGCTACCACATCCAAG

18S reverse: CCTCCAATGGATCCTCGTTA

*Dynlrb1* forward: CAACCTCATGCACAACTTCATC (exon 3)

*Dynlrb1* reverse: TCTGGATCACAATCAGGAAATAGTC (exon 4)

### RNA sequencing

Wild type and *Dynlrb1^+/-^* DRG ganglia were dissociated as described above and cells were plated on coverslips in F12 medium (ThermoFisher Scientific). Cells were washed in PBS at 12, 24 and 48 h after plating and total RNA was extracted using the Oligotex mRNA Mini Kit (Qiagen). RNA purity, integrity (RIN > 7) and concentration was determined. Three biological repeats were analyzed for each time point for each genotype.

RNA-sequencing libraries were prepared using the TrueSeq Stranded RNA kit (100ng). Libraries were indexed and sequenced by HiSeq4000 with 75 bp paired-end reads and at least 50M reads were obtained for each sample. Quality control (QC) was performed on base qualities and nucleotide composition of sequences, mismatch rate, mapping rate to the whole genome, repeats, chromosomes, key transcriptomic regions (exons, introns, UTRs, genes), insert sizes, AT/GC dropout, transcript coverage and GC bias to identify problems in library preparation or sequencing. Reads were aligned to the latest mouse mm10 reference genome (GRCm38.75) using the STAR spliced read aligner (ver 2.4.0). Average input read counts were 66.6M and average percentage of uniquely aligned reads were 86.3%. Total counts of read-fragments aligned to known gene regions within the mouse (mm10) ensembl (GRCm38.80) transcript reference annotation are used as the basis for quantification of gene expression. Fragment counts were derived using HTSeq program (ver 0.6.0). Genes with minimum of 5 counts for at least half of samples were selected. Filtered count data were normalized and examined for sequencing bias by PCA, and PC1 and PC2 of sequencing QC (described above) were regressed out. Differentially expressed transcripts were determined by Bioconductor package EdgeR (ver 3.14.0) at FDR <0.1. Scripts used in the RNA sequencing analyses are available at https://github.com/icnn/RNAseq-PIPELINE.git. Raw and processed data were deposited within the Gene Expression Omnibus (GEO) repository (www.ncbi.nlm.nih.gov/geo), accession number (GSE131455).

The gene list from the 48h time point was filtered according to the FDR (FDR<0.1, **Table S1**) and uploaded to the Ingenuity Pathway Analysis software suite (QIAGEN Bioinformatics). The 10 best high ranking pathways according to their score (- log(p-values)) were further evaluated and the WNT – β-catenin pathway selected for validation (**Table S2**).

### Intrathecal injection of AAV9 in mice

Intrathecal (IT) injections of adeno associated virus serotype 9 (AAV9) of 5 µl total volume were performed at the lumbar segment L4 using a sterile 10 µl Hamilton microsyringe fitted with a 30 gauge needle. For all constructs, we obtained titers in the range of 10^12-10^13 viral genomes/ml, which were used undiluted for intrathecal injections.

### Behavioral profiling

All assays were performed during the “dark” active phase of the diurnal cycle under dim illumination (∼10 lx); the ventilation system in the test rooms provided a ∼65 dB white noise background. Every daily session of testing started with a 2 h habituation period to the test rooms. A recovery period of at least 1 day was provided between the different behavioral assays.

### Home-cage locomotion test

Wild type and *RNX3-Dynlrb1*^-/-^ mice were monitored over 72 h using the InfraMot system (TSE Systems, Germany) in order to investigate possible alterations of basal motor activity and/or circadian rhythms. The spatial displacement of body-heat images (infra-red radiation) over time was tracked [59]. The first 24 h were considered as a habituation phase, while the mouse activity of following 48 h (2 consecutive dark/light cycles) was calculated. Statistical analysis was carried out with a one-way ANOVA followed by Tukey post-hoc multiple comparison test using the statistical software GraphPad Prism.

### Open field

The motility of wild type and *RNX3-Dynlrb1*^-/-^ mice was assessed in the open-field arena under 120lx [59].The total distance moved (cm), the time spent (wall, corner; s) and mean speeds (cm/s) were recorded using VideoMot2 (TSE System, Germany) or Ethovision XT11 (Noldus Information Technology, the Netherlands). The data were analyzed with the COLORcation software [60]. Statistical analysis was carried out with a one-way ANOVA followed by Tukey post-hoc multiple comparison test or two-way ANOVA (for speed and distance) using the statistical software GraphPad Prism.

### Accelerated rotarod

Wild type and *RNX3-Dynlrb1*^-/-^ mice and wild type mice intrathecally injected with AAV9-shControl or AAV9-shDynlrb1 were tested with the ROTOR-ROD system (83x91x61 - SD. Instruments, San Diego) for their balance/coordination ability [61, 62]. Mice were subjected to 3 trials, with 5 min inter-trial intervals. Rotarod acceleration was set at 20 rpm in 240 s. Latency to fall (s) was recorded and the average of the 3 trials was considered. Statistical analysis was carried out with two-way ANOVA using the statistical software GraphPad Prism.

### Catwalk Gait Analysis

Wild type and *RNX3-Dynlrb1*^-/-^ mice and wild type mice intrathecally injected with AAV9-shControl or AAV9-shDynlrb1 were assessed for their motor/balance coordination using the CatWalk as previously described [63]. Motivation was achieved by placing the home cage at the end of the runway. The test was repeated 3-5 times for each mouse. Data were collected and analyzed using the Catwalk Ethovision XT11 software (Noldus Information Technology, The Netherlands). The parameters reported for each animal are the paw stance, defined as the duration in sec of the contact of the paw with the glass surface, and the step sequence regularity index, defined as the number of normal step sequence patterns relative to the total number of paw placements. Statistical analysis was carried out with a one-way ANOVA followed by Tukey post-hoc multiple comparison test (Wild type and *RNX3-Dynlrb1*^-/-^ mice) or t-test (AAV injected mice) using the statistical software GraphPad Prism.

### Statistical analysis

Analysis of multiple groups was made using the ANOVA method. The choice between one- or two-way ANOVA was based on the requirements for identification of specific factors’ contribution to statistical differences between groups and were followed by the Tukey and the Sidak post hoc analysis tests respectively. For 2-groups analyses, unpaired Student’s t test was used. All analyses were performed using GraphPad Prism version 7.00 for MacOS (GraphPad Software, La Jolla, California, USA, https://www.graphpad.com/). The results are expressed as the mean ± standard error of the mean (SEM). All statistical parameters for specific analyses are reported in the figure legends of the paper. Statistically significant P-values are shown as *p < 0.05, **p < 0.01, ***p < 0.001 and ****p < 0.0001.

